# Spatial bias in GBIF data has limited impact on plant climate niche properties in Europe

**DOI:** 10.64898/2026.02.11.705286

**Authors:** Thibault Coquery, Erik Welk, Lotte Korell

## Abstract

**Aim:** The Global Biodiversity Information Facility (GBIF) is the most prominent source of species occurrence data for modeling climate niches, but exhibits strong unevenness in its data coverage across different geographic regions. The impact of this spatial bias on the reliability of GBIF-based plant climate niches in Europe remains unexplored. This study aims to address this gap, and to investigate whether the targeted integration of additional atlas data can reduce the potential impact of the spatial bias.

**Location:** Europe.

**Time period:** 1950s - 2024

**Major taxa studied:** European grassland plant species.

**Methods:** We analyzed the climate niches of a large number of grassland species, with diverse distribution patterns across Europe, based on a) GBIF and b) on an enriched version of GBIF with national atlas data from Eastern European countries (GBIF+), where data coverage is currently low in GBIF. We followed best practices in niche characterization, particularly by performing environmental subsampling. The accuracy in climate niche properties was determined by comparing niches based on GBIF and GBIF+ data with niches based on a careful implementation of expert range maps as reference dataset. We focused on niche optimum position and niche similarity. Additionally, we investigated how biogeographical indicators can predict variability in climate niche accuracy.

**Results:** Most species exhibited reliable climate niche characterization using GBIF data, especially for widely distributed species. Yet, reliability decreased with continentality; that is, when species were primarily distributed in Eastern Europe. Integrating additional data did not significantly reduce this bias in niche characterization.

**Main conclusions:** Despite the spatial bias in its records, GBIF can be used to reliably characterize the climate niches of many species in Europe if uneven sampling effort is accounted for. The laborious integration of additional data to address spatial bias does not yield the desired increase in niche reliability.

## Introduction

Accurate characterization of species realized climate niches is fundamental to contemporary ecological research. The realized niche describes the range of (climatic) conditions under which a species can maintain viable populations, derived empirically from the environmental conditions at known geographical occurrences (Hutchinson, 1957). Understanding these niches has become increasingly important for predicting species responses to climate change (Thuiller, Lavorel, & Araújo, 2005), informing conservation prioritization (Rathore & Sharma, 2023), and projecting future distribution shifts through species distribution models (SDMs; Franklin (2023)). As global environmental change accelerates, reliable climate niche estimates are essential for evidence-based conservation and management decisions. This is particularly critical for plant species, whose movements may not keep up with climate change (Corlett & Westcott, 2013).

The Global Biodiversity Information Facility (GBIF; https://www.gbif.org) has emerged as the dominant source of species occurrence data for climate niche modeling (Saran, Chaudhary, Singh, Tiwari, & Kumar, 2022). This international infrastructure hosts the world’s largest collection of species occurrence records, with an exponential increase in data available with time, in addition to being free and open-access (GBIF Secretariat, 2024). It aggregates datasets from museum collections, national atlases, research surveys and citizen science (Heberling, Miller, Noesgaard, Weingart, & Schigel, 2021). Consequently, the majority of recent niche modeling studies rely primarily or exclusively on GBIF data. However, this widespread dependence has raised concerns about spatial sampling bias inherent in these data (Troia & McManamay, 2016). Occurrence records in GBIF show pronounced geographic unevenness, with substantially higher sampling intensity and data mobilization in Western European countries compared to Eastern Europe and other regions (García-Roselló, González-Dacosta, & Lobo, 2025; Hughes et al., 2021; Meyer, Weigelt, & Kreft, 2016). This pattern reflects historical differences in scientific infrastructure, collection traditions and digitization efforts rather than underlying biodiversity patterns (Amano & Sutherland, 2013). More recent contributions of citizen science to global databases may even amplify the uneven data coverage in GBIF (Melis et al., 2025).

This spatial bias in GBIF data may directly affect the characterization of species’ climate niches. When occurrence records are geographically clustered, the resulting climate niche may disproportionately reflect environmental conditions from oversampled regions, thereby underrepresenting the full climatic range actually occupied by a species. For example, species whose distributions span both well- and poorly-sampled areas may appear to have narrower or truncated thermal tolerances, or shifted niche optima, compared to their true ecological breadth. This bias may be particularly pronounced for species whose ranges coincide with socio-economically driven sampling patterns that also correspond to major climatic gradients—such as the continentality gradient in Europe, where temperature seasonality and precipitation regimes vary sharply from oceanic western to continental eastern regions (Berg, Welk, & Jäger, 2017). Previous studies have documented that spatial sampling bias can alter niche breadth estimates, shift apparent niche centroids, and affect model predictions (Beck, Ballesteros-Mejia, Nagel, Kitching, & Ferrier, 2013; Beck, Böller, Erhardt, & Schwanghart, 2014; Hortal, Jiménez-Valverde, Gómez, Lobo, & Baselga, 2008). In contrast, other studies have revealed similar niche properties between GBIF data and other datasets less prone to spatial bias, suggesting that the magnitude of these effects can vary among taxa and geographic contexts (Alhajeri & Fourcade, 2019; Pender et al., 2019). The spatial bias for plants in Europe is well described, yet its potential impact on climate niche properties still needs to be investigated.

To address the spatial bias and the potential bias in climate niche estimates, several researchers have advocated for integrating multiple data sources beyond GBIF, particularly for regions with sparse coverage (Fletcher et al., 2019; Wetzel et al., 2018). Additional datasets might include national biodiversity databases, regional flora and fauna surveys (de Araujo, Quaresma, & Ramos, 2022). For years now dataset holders have been encouraged to mobilize their data in GBIF (Yesson et al., 2007), but still many valuable datasets are stored in different repositories which does not facilitate their compilation. However, a critical question remains largely unexplored: Does the often time-consuming integration of additional data sources actually produce more realistic climate niche estimates? Addressing this question requires an independent distribution dataset to serve as a reference for estimating niches and for comparing these estimates with those obtained from GBIF data alone and from GBIF data augmented with supplementary sources. In this context, expert-derived range maps offer a promising solution (Alhajeri & Fourcade, 2019; Hanberry, 2025; Rotenberry & Balasubramaniam, 2020). These maps, built upon comprehensive species knowledge, are less vulnerable than point occurrence data to sampling artifacts and niche truncation (Jetz, McPherson, & Guralnick, 2012).

In this study, we systematically compared the climate niches of plant species in Europe derived from different occurrence data sources against climate niches based on expert range maps as an independent reference. First, we quantified the potential bias in climate niche properties when using GBIF data and its inherent spatial bias. Then, following the recommended practice of addressing spatial bias by supplementing GBIF data with additional occurrence sources from four Eastern European countries, we evaluated whether the resulting climate niche characterization aligns better with those based on expert-derived distributions. We focused on European grassland species, which offer an ideal study system, as they represent an important habitat type that spans the full continentality gradient, and for which comprehensive expert range maps are well documented (Berg et al., 2017; Meusel & Jäger, 1965, 1978, 1992). To compare the niches based on the different datasets, we considered two important niche properties: niche optimum position and niche similarity, which takes niche size into account (Thuiller et al., 2005).

We expect (i) stronger biases in climate niche properties when using GBIF data for continental species—mainly distributed in Eastern Europe—than for oceanic species, which occur mostly in Western Europe; and (ii) that the climate niche properties estimated from the enriched GBIF dataset (GBIF+) will take intermediate values between those derived from GBIF alone and those from expert range maps, with larger shifts for continental than for oceanic species. Our findings aim to inform best practices for reliable climate niche modeling and to support evidence-based data compilation strategies, particularly when working with species distributed across regions of uneven sampling coverage.

## Methods

### Distribution datasets and species selection

We focused in this study on temperate grassland plants occurring in Eastern Europe, from Eurosiberian agricultural grasslands (*Molinio-Arrhenatheretea*, Tx. 1937) or natural dry grasslands (*Festuco-Brometea*, Br.-Bl. et Tx. ex Soó 1947). For each grassland species included in the analysis, we prepared three different distribution datasets. 1) The GBIF dataset was composed of the occurrence records available in GBIF until 2024 (https://www.gbif.org). As suggested in most studies, occurrence records were cleaned using tests from the ‘CoordinateCleaner’ package (Zizka et al., 2019). 2) The expert range map dataset consisted of point and polygon data, thoroughly drawn by hand based on the best available knowledge on species distributions (Meusel & Jäger, 1965, 1978, 1992). These maps distinguish main range, range islands and isolated occurrences where appropriate. They were recently digitized and made publicly available (Wesenberg et al., 2026). 3) The GBIF+ dataset was composed of the GBIF records from the first dataset, enriched with additional data from four Eastern European countries, based on national atlases which are not yet included in GBIF. These data from Poland, the Czech Republic, Hungary and Croatia are available online, though not as GIS files, but rather as map graphics of the country with point occurrence signatures in the recording grid cells where the species is reported (https://www.atlas-roslin.pl; https://pladias.cz/en; http://floraatlasz.uni-sopron.hu; https://hirc.botanic.hr). To extract these data, we first georeferenced the respective graphic files. Then, we vectorized the data semi-automatically, based on the respective color values and spatial pattern of the respective target raster cells. Finally, we pooled these additional data with the GBIF dataset (Fletcher et al., 2019).

For each species, we considered its indicator value for continentality (C-value), following Berg et al. (2017). These biogeographic indicator values, revised from Ellenberg’s original concept reflect both a species’ climatic requirements and its position along the Eurasian oceanity–continentality gradient. C-values range from 1 for the most oceanic species, typically distributed in the westernmost parts of Europe, to 10 for the most continental species, primarily occurring in Central Asian regions. In Eastern Europe, species with different biogeographic affinities co-occur: oceanic species reach their eastern distributional limits, intermediate species have their core ranges, and continental species extend their western limits there. In contrast, some species are widely distributed across Eurasia, spanning both Western regions with dense GBIF coverage and Eastern regions with sparser data. Because these species are not clearly associated with either side of the oceanity–continentality gradient, they were treated as indifferent to continentality and were not assigned a C-value. In this study we included species for which we could compile the three distribution datasets and for which the indicator value for continentality was revised in Berg et al., 2017. Nomenclature and taxonomy follow *Rothmaler – Exkursionsflora von Deutschland* (Müller, Ritz, Welk, & Wesche, 2021).

### Climate niche characterization

Climate niche properties were quantified by summarizing the climatic space occupied by each species, using the values of selected bioclimatic variables extracted at their occurrence locations. To this end, we re-sampled the distribution datasets using a 0.1° stratification raster (approximately 11 km at the equator) over Europe, which was delimited to the west by Iceland, to the east by the Ural Mountains and to the south by Gibraltar (25° W, 72°N, 60°E, 36°S). For each dataset, we sampled one occurrence point per raster cell containing at least one occurrence record or overlapped with a polygon of the expert range maps. This rasterization step reduced the overrepresentation of certain areas in Western Europe in GBIF to some extent. At all sampled point positions, we then extracted the values of the 19 BioClim variables (climatology from 1980 to 2010; Karger et al. (2017)).

Although expert range maps typically have a low omission rate (i.e., few true occurrences missed), they are well known to have a high commission rate due to their coarse spatial resolution (Jetz et al., 2012; Mainali, Hefley, Ries, & Fagan, 2020). For example, they often include entire mountain ranges due to the original map scale and cannot precisely exclude areas at very high altitudes. These commission errors can introduce bias into estimates of the species’ climate niche (Zhang et al., 2025). We developed an approach to systematically remove unsuitable grid cells, which are often associated with extreme climatic values. For each species, we plotted mean annual temperature and log-transformed precipitation values of raster cells within the expert range map. Outliers typically appeared as isolated points far from the dense cluster of data, representing cells with very low or very high temperature or precipitation. To remove these outliers, we fitted a convex hull around the points and iteratively removed its outer vertices—ten times for species with extensive niches and fewer iterations for species with smaller data ranges. With this *convex hull peeling process*, most of the outlier points in the scatterplot were removed (Fig. S1). Plotting these occurrences back on a map of Europe, confirmed them as areas of expected potential unsuitable climate conditions. We then performed climate niche analyses based on expert range maps using the “cleaned” dataset that remained after the peeling process.

Since we were interested in comparing the climate niches between data-sources, rather than between species, we defined species-specific climate spaces using the first two components of a principal component analysis (PCA; Broennimann et al. (2012)). For each species, we only considered the climate variables that were not highly intercorrelated (Pearson’s correlation coefficient r < 0.7). We calibrated the PCA multidimensional climate space using the remaining climate variables and the three distribution datasets combined. Next, we performed eco-geographical subsampling of the grid cells for each of the three datasets based on their PCA scores for the first two components, using the ‘occfilt_env’ command from the ‘flexsdm’ package with nbins = 25 (Velazco, Rose, de Andrade, Minoli, & Franklin, 2022). This step ensured a homogeneous density of occurrences within the climate niche and prevented overrepresentation of climate conditions in areas with dense spatial coverage (Fig. S2; Varela, Anderson, García-Valdés, and Fernández-González (2014)). The filtered occurrences for each dataset, delimited by a minimum convex hull polygon in the two-dimensional PCA space, defined the respective climate niche (Beck et al., 2013).

### Comparison of climate niches

For each species, we compared three climate niches derived from different distribution datasets: GBIF, GBIF+, and expert range maps. The niche based on expert range maps served as an independent reference because this dataset is expected to have the lowest spatial bias and reduced commission error (Jetz et al., 2012). We first compared the GBIF-based niche with the expert range map to evaluate the potential influence of spatial bias in GBIF records. Next, we repeated the comparison using the enriched GBIF+ dataset to assess whether the additional data reduced this bias. The first niche property we examined was the position of the niche optimum. We defined the centroid of each niche as the median of filtered occurrence points. Bias in niche position was quantified as distance between the GBIF or GBIF+ centroid and that of the expert range map (Perez-Navarro et al., 2022). Because the first axis of the climate space explained more variance than the second, we used the Mahalanobis distance to calculate differences. The relative change in Mahalanobis distance between GBIF and GBIF+ indicated whether enrichment reduced bias—a negative change meaning that the GBIF+ centroid was closer to the expert-based centroid. We also quantified similarity in overall niche shape and extent using Sørensen’s similarity index, which measures the proportion of shared climatic conditions and accounts for differences in niche size (Godsoe & Case, 2015). Index values close to 1 indicate high similarity and low bias. The relative change in Sørensen’s index between GBIF and GBIF+ provided an additional measure of bias reduction—positive values indicating increased similarity with the expert-based niche.

The statistical analyses were conducted in R 4.5.0 (R Core Team, 2025). In a first step, we analyzed potential biases in niche properties using the GBIF dataset. We examined the distribution of the two metrics (see above) among all the species included in the study. Then, we compared the metrics between widely distributed species indifferent to continentality and non-indifferent species for which a continentality indicator value could be assigned. Finally, for the latter group of species, we evaluated whether species with higher continentality values exhibited greater bias in both niche properties. To do so, we considered the discrete C-values from 1 to 10 as a continuous variable because they represent an eco-geographical gradient. Treating the centroid distance as response variable, we performed a Gamma regression, and for Sørensen’s index a Beta regression. In a second step, we analysed the relative changes in centroid distance and niche similarity between the GBIF and GBIF+ datasets, still using the expert range–derived niches as the reference. We examined the distributions of these relative changes and compared the patterns of Mahalanobis distance and Sørensen’s index between species that were indifferent and those that were restricted in continentality. Within the group of continentality restricted species, we further investigated the influence of their C-values on these metrics. As a conservative approach, we excluded half of the species whose GBIF-based niches showed the smallest initial bias, as minor positional changes between already similar centroids can produce disproportionately high relative changes. Where necessary, response variables were transformed to meet the assumptions of linear and generalized linear models.

## Results

A total of 175 species that met our selection criteria were included in the analysis. Of these species, 69 were indifferent to continentality, while 106 had a continentality indicator value assigned based on their restricted range along the gradient (Table S1). These C-values ranged from 3 to 8, and covered the eco-geographical gradient of species distributions which occur in Eastern Europe (Fig. 1). Depending on the species, the PCA climate space explained between 51.5 % and 73.8 % of the respective data variance.

**Figure 1.**
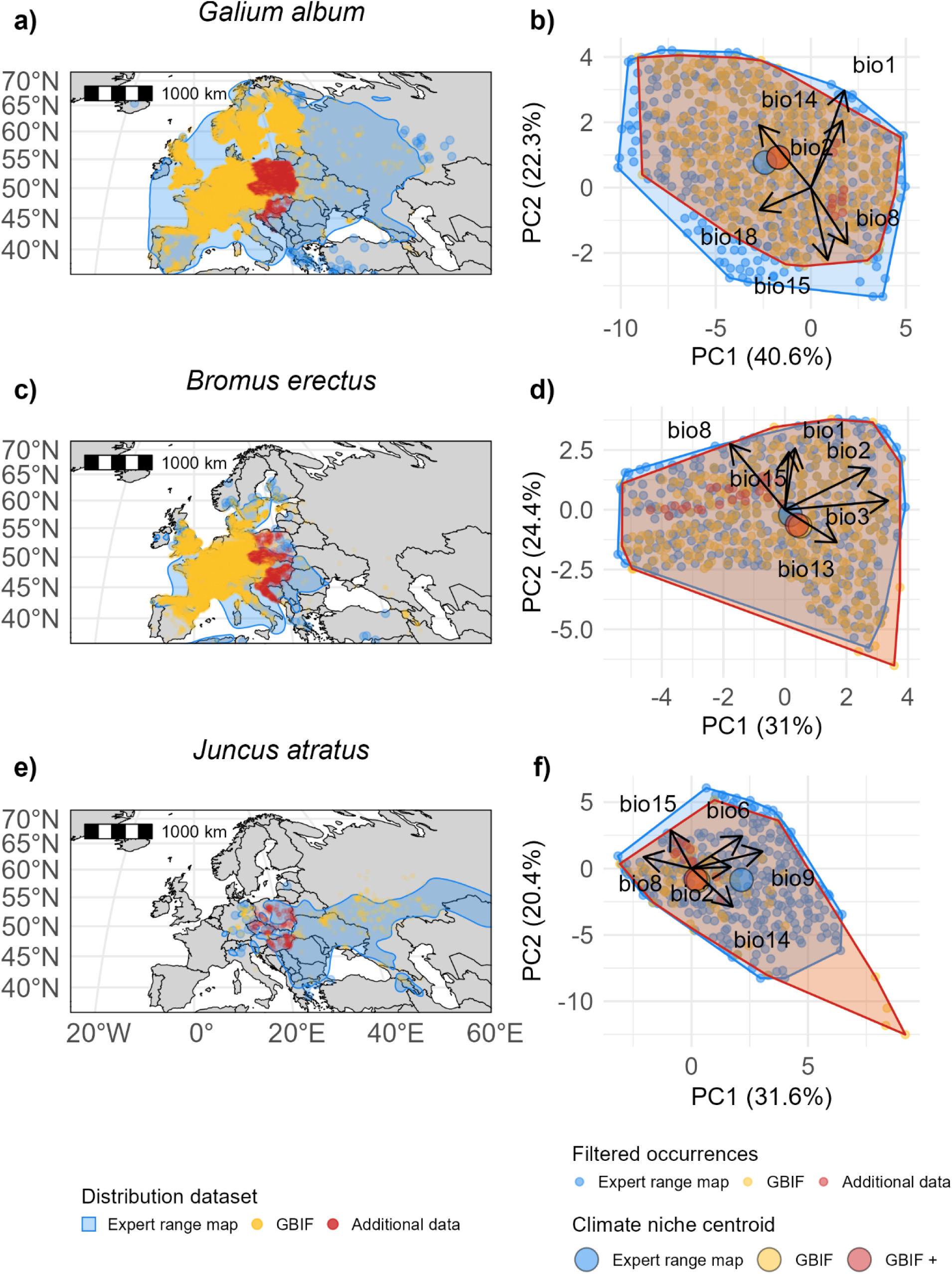
Examples of three grassland plant species included in the analysis with their distribution datasets in Europe (left), used to fit climate niches (right). The three datasets are: expert range map, GBIF data, and GBIF enriched with national atlas data from Poland, the Czech Republic, Hungary and Croatia (GBIF+). Climate niches are defined in a two-dimensional PCA climate space after outlier removal in the expert range maps and environmental subsampling, using BioClim variables. Niche edge is delimited by the minimum convex hull polygon of the filtered occurrences. Note that the niches based on GBIF (yellow) perfectly overlap with niche based on GBIF+ (red). Niche centroid, defined as the median point, is shown, as well as the deviance explained by PC1 and PC2. *Galium album* (top row) is an example of a species indifferent to continentality, *Bromus erectus* (middle row) of a species mostly distributed in Western Europe (C-value of 4) and *Juncus atratus* of a species mostly distributed in Eastern Europe (C-value of 8). Maps with Mollweide projection.

### Potential bias in GBIF-derived climate niche properties

When we compared the niche based on GBIF with the niche based on the expert range map for each species, we found low differences for both considered niche properties and a similar impact of the predictors describing the supposed trend in bias impact (Fig. 2). First, the niche optimum position was very similar between the GBIF and expert range datasets for most species, with a distance between niche centroids smaller than 0.1 (Fig. 2a). 90% of the species had a distance between centroids less than 0.7. For some species, however, the bias was greater, e.g., reaching a maximal distance of 4.5 for *Cardamine parviflora*. Specifically, the bias in niche optimum position was lower for species indifferent to continentality than for non-indifferent species (e = −1.14, SE = 0.27, t = −4.28, P < 0.001, Fig. 2b). Both groups had a similar low median value (respectively 0.04 and 0.13), but species indifferent to continentality had smaller standard deviation (0.14 versus 0.83). Among non-indifferent species, the greater the continentality indicator value was, the greater was the distance between centroids (e = 0.56, SE = 0.10, t = 5.35, P < 0.001, Fig. 2c). Regarding niche similarity, many species had high values of Sørensen’s similarity index between niches derived from the GBIF dataset and the expert data-based niches (Fig. 2d). The smallest value was 0.55 for *Carex supina*. Species indifferent to continentality showed higher similarity index values than non-indifferent species (e = 0.43, SE = 0.09, z = 5.04, P < 0.001, Fig. 2e). For the latter group, Sørensen’s index values decreased with increasing C-values (e = −0.16, SE = 0.04, z = −4.23, P < 0.001, Fig. 2e), indicating a greater bias in niche similarity for more continental species. Distance between centroids and Sørensen’s index values were negatively correlated, meaning that species with a large niche optimum position bias also showed a large niche similarity bias (Pearson’s correlation coefficient r = - 0.70).

**Figure 2.**
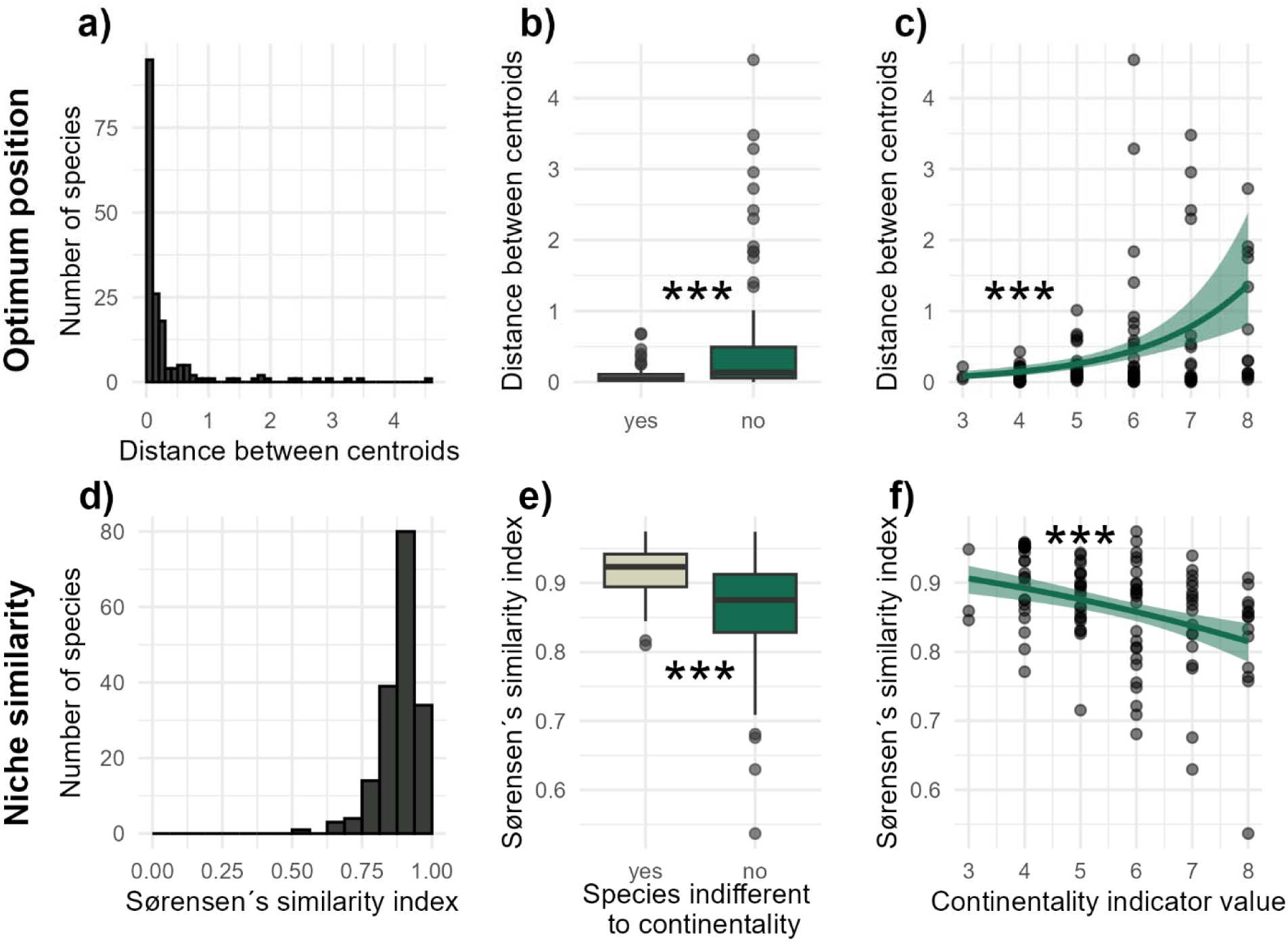
Bias in climate niche properties of European grassland species using GBIF. Reference niche for comparison with GBIF was based on cleaned expert range maps. Bias in optimum position was quantified with the Mahalanobis distance between the centroids from each of the two niches (top row) and niche similarity was quantified with Sørensen’s index (bottom row). Left column shows the distribution of the two metrics for the 175 considered species. Middle column shows differences between species indifferent (yes) and not-indifferent to continentality (no). Boxplots with the median (horizontal line), interquartile range (IQR; box), and the 1.5 × IQR whiskers. Points represent individual species. Right column shows the impact of the continentality indicator value (C-value), with model mean prediction and 95% CI (n = 106). Linear model with ln-transformation was used in (b), Gamma regression in (c) and Beta regression in (e) and (f). *** indicates P < 0.001.

### Impact of additional data on climate niche properties (GBIF+)

Since for many species the niche centroids were highly similar (see above), we evaluated the impact of the additional data on the niche optimum position for only half of the species with larger distance between centroids (distance > 0.1; GBIF vs. expert range map data). Even for the majority of these considered species, the additional data only marginally changed or did not change the centroid position at all (Fig. 1b, Fig. 3a). For some species, the relative change in distance between GBIF+ and GBIF was negative; that is, the GBIF+ centroid was closer to the reference centroid than the GBIF centroid (Fig. 1d). For some other species, the additional data shifted the centroid position further away from the reference centroid than it was with GBIF dataset, i.e., the relative change in distance was positive (Fig. 1f). The mean relative change in distance was −3.2%. Since most of the considered species were not indifferent to continentality, we did not compare the relative change in distance with species that were indifferent to continentality. Against expectations, the continentality indicator values did not explain patterns of relative change in distance between centroids (e = −0.10, SE = 0.19, t = - 0.52, P = 0.607, Fig. 3b). Finally, the additional data also had a very limited impact on niche similarity: 148 out of 175 species showed no difference in Sørensen’s similarity index (Fig. 3c). The index value decreased for nine species and increased for 18 species, meaning that GBIF+ niche was more similar to the niche based on the expert range map than the GBIF niche. Due to the limited impact of the additional data on niche similarity change, we did not investigate the effect of the predictors describing species distribution patterns.

**Figure 3.**
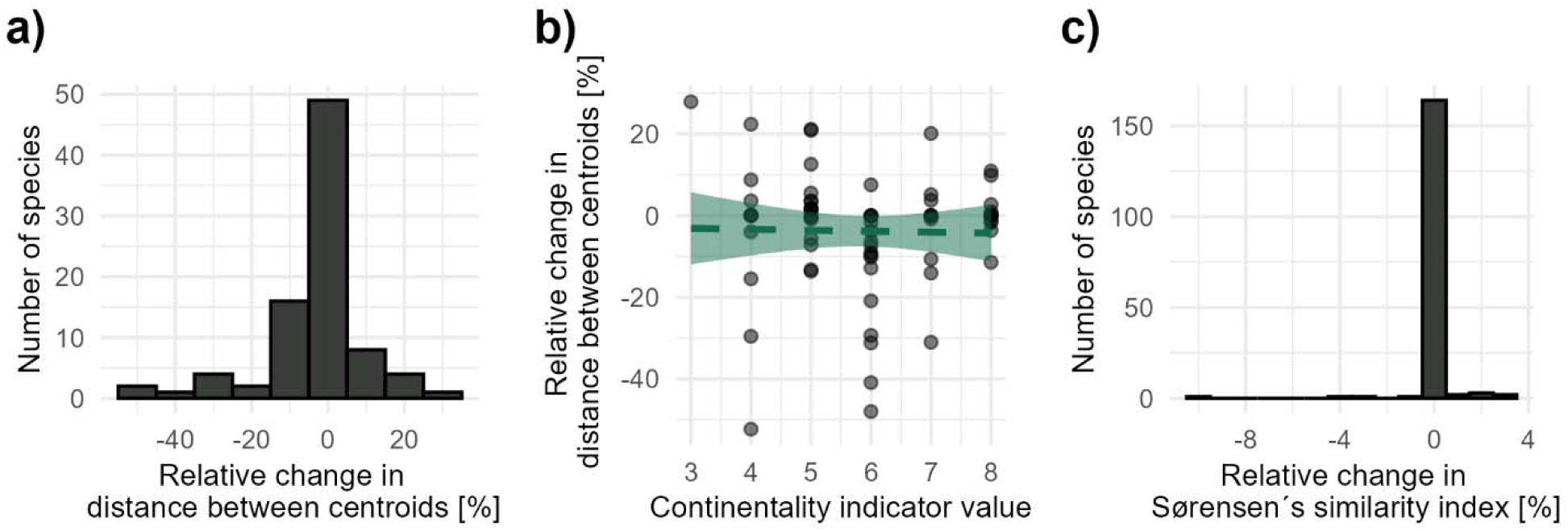
Impact of additional national atlas data from East European countries integrated to GBIF on the bias in climate niche properties. Relative change in bias is based on GBIF+ niche compared to GBIF niche, with in both cases the niche based on cleaned expert range map as reference. Change in optimum position is quantified by change in Mahalanobis distance between centroids (a, b) and change in niche similarity is quantified by change in Sørensen’s index (c). Change in optimum position was not assessed for species with low bias with GBIF (upper half of the dataset considered, n = 87). In b), mean prediction and 95% CI of the linear model (signed log-transformation) are shown.

## Discussion

This study demonstrates that GBIF is a reliable source of distribution data for accurately characterizing the realized climatic niches of many European plant species. The limited effect of spatial sampling bias on estimating climatic niche properties supports the use of carefully cleaned GBIF data for climate niche modelling and helps put concerns about such bias into perspective. These findings also validate previous studies that modelled the climatic niches of European plants using GBIF-derived data when appropriate cleaning and subsampling strategies were applied. However, as expected, species with predominantly continental distributions exhibited greater bias in climatic niche characterization. Such bias may influence downstream analyses, including projections of species distributions under climate change. Moreover, the labor-intensive integration of additional data sources to compensate for lower data coverage in parts of Eastern Europe did not lead to the anticipated improvement in the accuracy of climatic niche estimates.

Our main finding aligns with studies testing a similar question but for different taxa and spatial extents (e.g., rodents, worldwide: Alhajeri and Fourcade (2019); avian order Galliformes, worldwide: Rotenberry and Balasubramaniam (2020); *Carex* genus, North America: Pender et al. (2019)). GBIF data exhibit clear spatial biases across Europe, with dense coverage in Western countries and sparse records in the East. Nevertheless, even in low-coverage regions such as Ukraine and Russia, some GBIF occurrences fall within species’ distribution ranges (Fig. 1a,e). Many of these records originate from citizen-science platforms such as iNaturalist rather than national databases (Ivanova & Shashkov, 2020). Despite their limited number, these occurrences are still informative, as they represent the dominant climatic conditions of these largely homogeneous, continental lowland regions. In contrast, countries like France and Germany feature considerable environmental heterogeneity over relatively small spatial scales, but exhibit higher GBIF coverage (Meyer, Kreft, Guralnick, & Jetz, 2015). Consequently, while data density varies geographically, the strong climatic diversity captured in Western Europe may already encompass much of the European climatic niche space. This pattern likely reduces the overall impact of the east–west spatial imbalance in GBIF coverage on estimated climate niche properties, especially when seen in combination with environmental subsampling strategies.

Although the overall bias in GBIF data was generally low for grassland species and for both niche properties examined, clear differences emerged among species with different European distribution patterns. Species indifferent to continentality—those broadly distributed across Europe and spanning areas of contrasting data coverage—showed lower bias in niche properties than species with more geographically or climatically restricted ranges (Fig. 2b,e). These widespread species also exhibited broader niche breadths. Because their large ranges encompass a wide spectrum of climatic conditions and range coverage can be low (Meyer, Jetz, Guralnick, Fritz, & Kreft, 2016), it might be assumed that GBIF records miss certain parts of their environmental variation. However, even if a specific climatic condition is underrepresented in one area, similar conditions elsewhere in their range are often sampled, reducing bias overall. Among species sensitive to continentality, those with low continentality values—mostly occurring in well-sampled Western European regions—also had relatively low GBIF biases. Conversely, species associated with higher continentality values showed greater niche property bias (Fig. 2c,f). Eastwards, with increasing continentality, the chance of a species’ range being located in under-sampled regions rises, particularly for species with restricted distributions. These findings suggest that the effect of GBIF’s spatial bias depends strongly on species’ range size and climatic associations. Caution is therefore warranted when modelling the climate niches of continental species using uncorrected GBIF data (Bystriakova, Peregrym, Erkens, Bezsmertna, & Schneider, 2012).

Since for most of the species included in the analysis the bias in the climate niche properties was very low in GBIF, the time investment required to integrate additional data to achieve more accurate niches would not have been worthwhile. Further, when the bias was rather large, the additional data often did not lead to more reliable climate niches. Contrarily to our expectations, the improvement in niche characterization was not greater for continental species. Hence, the data from Poland, the Czech Republic, Hungary and Croatia did not provide new climatic conditions that had not been already covered in GBIF (Štípková, Tsiftsis, & Kindlmann, 2024). This investment may have been worthwhile with additional data from more eastern countries, such as Ukraine, Belarus or Oblasts of European Russia, where data coverage in GBIF is even sparser. However, floristic atlas data for these regions are unavailable or too coarse (Dudov & Seregin, 2025). This further emphasizes the importance of citizen science data integrated into GBIF in countries representative of very continental climate conditions (Dylewski et al., 2025; Seregin et al., 2020).

Our conclusions are based on the use of expert range maps as a reference dataset. Despite the methodological difference between GBIF (i.e., point occurrence data) and expert range maps (i.e., polygon data, drawn by hand based on expert knowledge) these two very different approaches to distribution data compilation yielded consistent climate niche properties (Alhajeri & Fourcade, 2019; Rotenberry & Balasubramaniam, 2020). This consistency also speaks for the relevance of the expert range map as an independent dataset (Hanberry, 2025). For the expert range maps, the polygons were drawn based on an in-depth review of literature records based on herbarium specimens that were likely digitized and included in GBIF. A difference between the two datasets is that GBIF contains more recent data than the expert range maps, which were compiled between the 1950s and 1980s. This similarity furthermore supports the *convex hull peeling process* that we developed to remove false positives from expert range maps (Fig. S1). This procedure could be applied to other studies using expert range maps for climate niche modeling to address this issue and enhance model accuracy (Zhang et al., 2025). Our study contributes to the growing body of literature emphasizing the importance of expert range maps in macroecological studies (Aronsson, Zizka, Antonelli, & Faurby, 2024; Oeser et al., 2024; Xiao et al., 2025).

This study showed that reliable niche characterization can be achieved using GBIF, despite its inherent spatial bias. However, we would like to emphasize the importance of environmental subsampling (Pili, Leroy, & Zurell, 2025). Without subsampling, the niche optimum position is driven toward the climatic conditions most represented in the dataset, i.e., those from areas with the best GBIF coverage (Fig. S2). Therefore, environmental subsampling reduces the potential impact of spatial bias on accurate niche characterization and should be a *sine qua non* step in niche modeling workflows (Varela et al., 2014). Finally, we acknowledge that some species may differ in their spatial bias across Europe (Meyer, Jetz, et al., 2016). Conspicuous species, species of public interest, and species of conservation concern may be less likely to be underrepresented in areas with less coverage (Tyler et al., 2012), and are therefore less likely to introduce bias in the characterization of their climate niches. Nevertheless, we included a large number of species in the analysis to represent the taxonomic and distributional diversity of grassland species in Eastern Europe, regardless of potential species-specific differences in spatial bias.

The heterogeneous sampling effort between Western and Eastern Europe is not limited to plants; rather it has been described across a variety of taxa (García-Roselló, González-Dacosta, & Lobo, 2025). Hence, our findings may apply to other taxa in Europe, not just grassland plant species. This may be particularly true for other taxa that, like vascular plants, are relatively well represented in GBIF, such as birds and mammals (Troudet, Grandcolas, Blin, Vignes-Lebbe, & Legendre, 2017). Patchy and clustered occurrences may not be such an issue if the entire range of suitable climate conditions for the considered species is represented in GBIF occurrences, even if there are just a few records, particularly in Eastern Europe. However, this may not be the case for taxa that are less well represented in GBIF, especially invertebrates (Beck et al., 2014; García-Roselló, González-Dacosta, & Lobo, 2023; Hughes et al., 2021).

## Conclusions

In summary, despite uneven geographic coverage across Europe, GBIF data can provide reliable estimates of plant climatic niche properties when overrepresented climatic conditions are explicitly accounted for. However, for some species, the risk of bias remains substantial, so researchers should carefully assess which taxa data are most vulnerable (e.g., those with more continental distributions) rather than relying on GBIF data alone. Integrating multiple data sources can improve sampling completeness, but its effect on climatic niche accuracy may be modest and context-dependent, especially considering the effort required. Such integration is most valuable when it adds poorly represented climatic conditions rather than merely filling geographic gaps. A key challenge arises when certain climates are absent from both GBIF and complementary databases. To address this, we advocate greater support for citizen science initiatives, particularly in underrepresented regions, as these can enhance data coverage while remaining compatible with bias-correction approaches (Rocchini et al., 2023).

## Author Contributions

Conceptualization: T.C., E.W., L.K.; Data provision: E.W.; Data analysis and visualization: T.C.; Writing – original draft: T.C.; writing – review and editing: all authors; supervision and funding acquisition: L.K.

## Acknowledgements

We thank the numerous citizen scientists and institutions who contributed occurrence records to GBIF.

## Data and Code Availability Statement

Data and R code used to conduct the analyses will be made available in Dryad upon acceptance of the manuscript: https://doi.org/10.5061/dryad.9ghx3ffzf; Reviewer link: http://datadryad.org/share/LINK_NOT_FOR_PUBLICATION/VsN2R_QICaEJFdulB6O7gXz5oGTYYCf57jocCQPfyZI

