## Supplementary material for "Spatial bias in GBIF data has limited impact on plant climate niche properties in Europe": Table S1

**Table S1**. **List of the 175 European plant grassland species included in the analysis.** The C-values refer to the revised indicator values for continentality of Ellenberg (Berg et al. 2017). These values range from 1 to 10, in this species list from 3 to 8. “x” means that the species is indifferent to continentality (n = 69).

| **Species** | **C-value** |
| --- | --- |
| *Achillea millefolium* | x |
| *Achillea pannonica* | 6 |
| *Achillea ptarmica* | x |
| *Achillea setacea* | 8 |
| *Adenophora liliifolia* | 7 |
| *Adonis vernalis* | 8 |
| *Agrimonia eupatoria* | x |
| *Agrostis canina* | x |
| *Ajuga genevensis* | 5 |
| *Allium angulosum* | 7 |
| *Allium oleraceum* | x |
| *Alopecurus myosuroides* | x |
| *Alopecurus pratensis* | x |
| *Angelica palustris* | 7 |
| *Angelica sylvestris* | x |
| *Anthemis tinctoria* | 6 |
| *Anthoxanthum odoratum* | x |
| *Anthriscus sylvestris* | x |
| *Anthyllis vulneraria* | x |
| *Arabis hirsuta* | 4 |
| *Arrhenatherum elatius* | 4 |
| *Asperula cynanchica* | 5 |
| *Aster amellus* | 7 |
| *Astragalus exscapus* | 8 |
| *Bellis perennis* | 4 |
| *Blysmus compressus* | 6 |
| *Brachypodium pinnatum* | x |
| *Briza media* | 4 |
| *Bromus erectus* | 4 |
| *Bromus sterilis* | x |
| *Campanula cervicaria* | 6 |
| *Campanula rotundifolia* | x |
| *Cardamine parviflora* | 6 |
| *Carex buxbaumii* | 7 |
| *Carex distans* | x |
| *Carex humilis* | 6 |
| *Carex supina* | 8 |
| *Centaurea jacea* | 6 |
| *Cichorium intybus* | x |
| *Cirsium arvense* | x |
| *Cirsium oleraceum* | 5 |
| *Cirsium palustre* | x |
| *Clinopodium vulgare* | x |
| *Cnidium dubium* | 7 |
| *Colchicum autumnale* | 4 |
| *Coronilla vaginalis* | 4 |
| *Crepis capillaris* | 4 |
| *Crepis mollis* | 5 |
| *Cynosurus cristatus* | 4 |
| *Daucus carota* | x |
| *Dianthus carthusianorum* | 5 |
| *Echium vulgare* | x |
| *Epilobium hirsutum* | x |
| *Equisetum ramosissimum* | 7 |
| *Eryngium campestre* | x |
| *Erysimum crepidifolium* | 6 |
| *Euphorbia angulata* | 4 |
| *Euphorbia cyparissias* | x |
| *Euphorbia palustris* | 6 |
| *Euphorbia seguieriana* | 7 |
| *Falcaria vulgaris* | 7 |
| *Festuca valesiaca* | 8 |
| *Filipendula ulmaria* | x |
| *Fumana procumbens* | 5 |
| *Galium album* | x |
| *Galium boreale* | 6 |
| *Galium verum* | x |
| *Genista pilosa* | 4 |
| *Gentiana asclepiadea* | 5 |
| *Geranium molle* | 4 |
| *Geranium palustre* | 5 |
| *Geranium pyrenaicum* | 4 |
| *Geranium sylvaticum* | 6 |
| *Geum rivale* | x |
| *Gladiolus palustris* | 6 |
| *Gratiola officinalis* | 6 |
| *Helianthemum canum* | 5 |
| *Heracleum sphondylium* | x |
| *Hippocrepis comosa* | 3 |
| *Holcus lanatus* | 4 |
| *Hornungia petraea* | 5 |
| *Hypericum elegans* | 7 |
| *Hypericum perforatum* | x |
| *Hypericum tetrapterum* | 4 |
| *Inula britannica* | 7 |
| *Iris sibirica* | 6 |
| *Iris spuria* | 6 |
| *Juncus acutiflorus* | x |
| *Juncus atratus* | 8 |
| *Juncus compressus* | x |
| *Juncus conglomeratus* | x |
| *Juncus effusus* | x |
| *Juncus inflexus* | x |
| *Juncus tenuis* | 6 |
| *Knautia arvensis* | x |
| *Koeleria macrantha* | x |
| *Lathyrus pratensis* | x |
| *Leontodon hispidus* | x |
| *Leucanthemum vulgare* | x |
| *Linum catharticum* | x |
| *Linum tenuifolium* | 6 |
| *Lolium perenne* | x |
| *Lotus corniculatus* | x |
| *Lotus pedunculatus* | 4 |
| *Lysimachia nummularia* | x |
| *Lysimachia vulgaris* | x |
| *Lythrum salicaria* | x |
| *Malva moschata* | 4 |
| *Medicago falcata* | x |
| *Medicago lupulina* | x |
| *Melilotus officinalis* | 8 |
| *Onobrychis arenaria* | 8 |
| *Ophioglossum vulgatum* | x |
| *Oxytropis pilosa* | 8 |
| *Pastinaca sativa* | x |
| *Phleum phleoides* | 6 |
| *Phyteuma orbiculare* | 5 |
| *Pimpinella saxifraga* | x |
| *Plantago lanceolata* | x |
| *Plantago major* | x |
| *Plantago media* | 7 |
| *Poa palustris* | 7 |
| *Poa pratensis* | x |
| *Poa trivialis* | x |
| *Polemonium caeruleum* | 7 |
| *Polygala amarella* | 5 |
| *Polygala comosa* | 6 |
| *Potentilla anserina* | x |
| *Potentilla heptaphylla* | 6 |
| *Primula veris* | x |
| *Prunella grandiflora* | 5 |
| *Pulicaria dysenterica* | x |
| *Pulsatilla vulgaris* | 5 |
| *Ranunculus aconitifolius* | 4 |
| *Ranunculus bulbosus* | 4 |
| *Ranunculus illyricus* | 8 |
| *Ranunculus repens* | x |
| *Rhinanthus alectorolophus* | 3 |
| *Rhinanthus minor* | x |
| *Rumex thyrsiflorus* | 7 |
| *Salvia pratensis* | 5 |
| *Saxifraga granulata* | 4 |
| *Scabiosa canescens* | 6 |
| *Scabiosa ochroleuca* | 8 |
| *Scorzonera humilis* | 5 |
| *Scorzonera purpurea* | 8 |
| *Scutellaria hastifolia* | 8 |
| *Selinum carvifolia* | 5 |
| *Silaum silaus* | 3 |
| *Sonchus oleraceus* | 4 |
| *Stachys recta* | 6 |
| *Stellaria graminea* | x |
| *Stipa capillata* | 7 |
| *Succisa pratensis* | x |
| *Succisella inflexa* | 6 |
| *Symphytum officinale* | x |
| *Tetragonolobus maritimus* | 5 |
| *Teucrium montanum* | 5 |
| *Thalictrum flavum* | x |
| *Thalictrum lucidum* | 6 |
| *Thesium linophyllon* | 5 |
| *Trifolium dubium* | 4 |
| *Trifolium fragiferum* | x |
| *Trifolium montanum* | 5 |
| *Trifolium repens* | x |
| *Trifolium spadiceum* | 5 |
| *Trinia glauca* | 5 |
| *Trisetum flavescens* | 4 |
| *Valerianella carinata* | x |
| *Veronica arvensis* | x |
| *Veronica prostrata* | 8 |
| *Vicia cracca* | 7 |
| *Vicia sepium* | x |
| *Viola elatior* | 7 |
| *Viola kitaibeliana* | 6 |
