## Supplementary material for "Spatial bias in GBIF data has limited impact on plant climate niche properties in Europe": Fig. S1

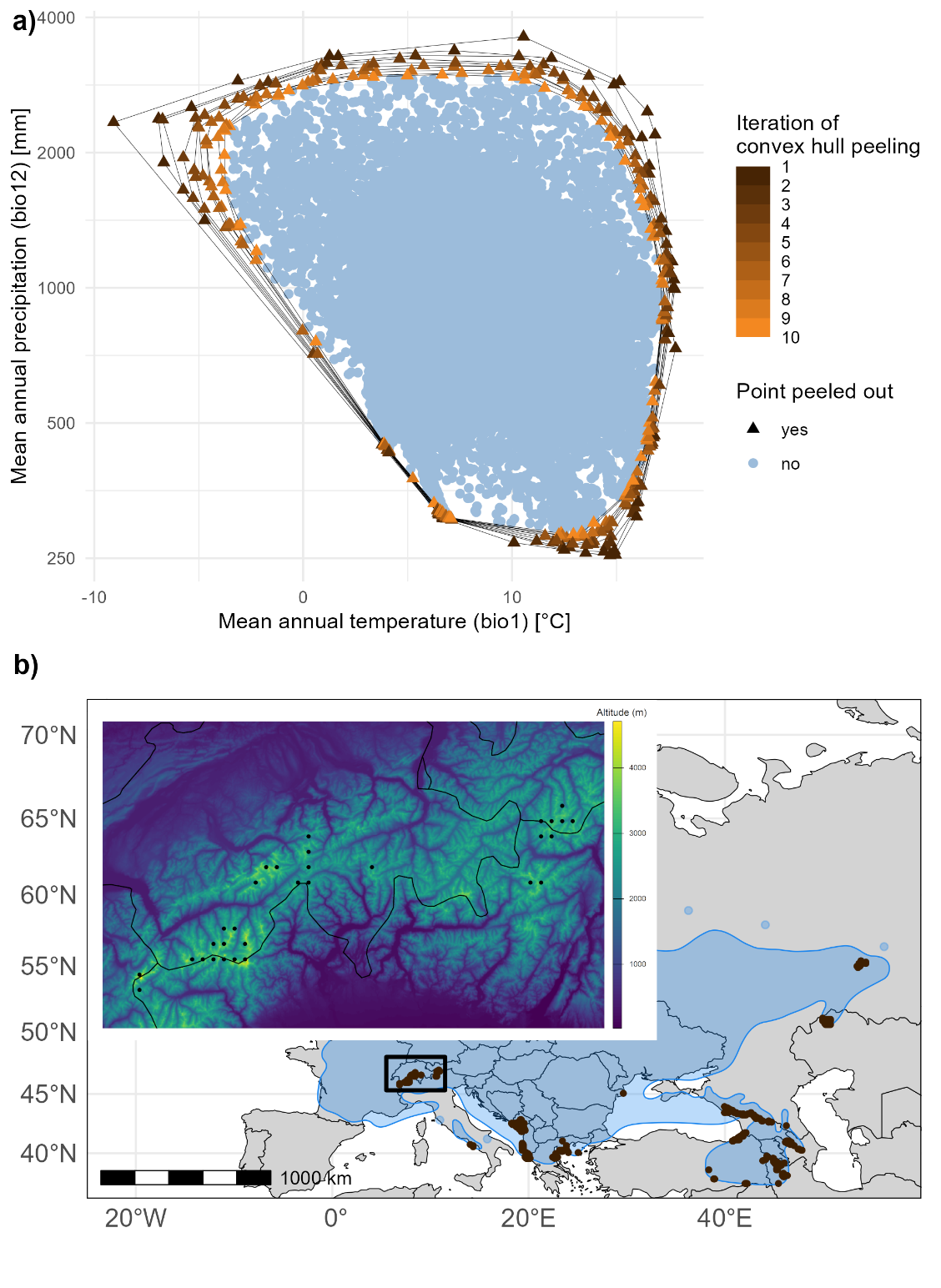

**Figure S1. Convex peeling process to remove potential false positive values in expert range maps with the example of *Ajuga genevensis*.** a) Removal of outlier points of expert range map in a scatterplot with mean annual temperature and mean annual precipitation (ln-transformed) on the axes. Outliers are likely vertices of convex hulls which are successively removed; b) Geographic position of removed points (black) in the expert range map (blue). A zoom on the Alps as insert is shown with altitude (retrieved from the ‘elevatr’ package).
