## Supplementary material for "Spatial bias in GBIF data has limited impact on plant climate niche properties in Europe": Fig. S2

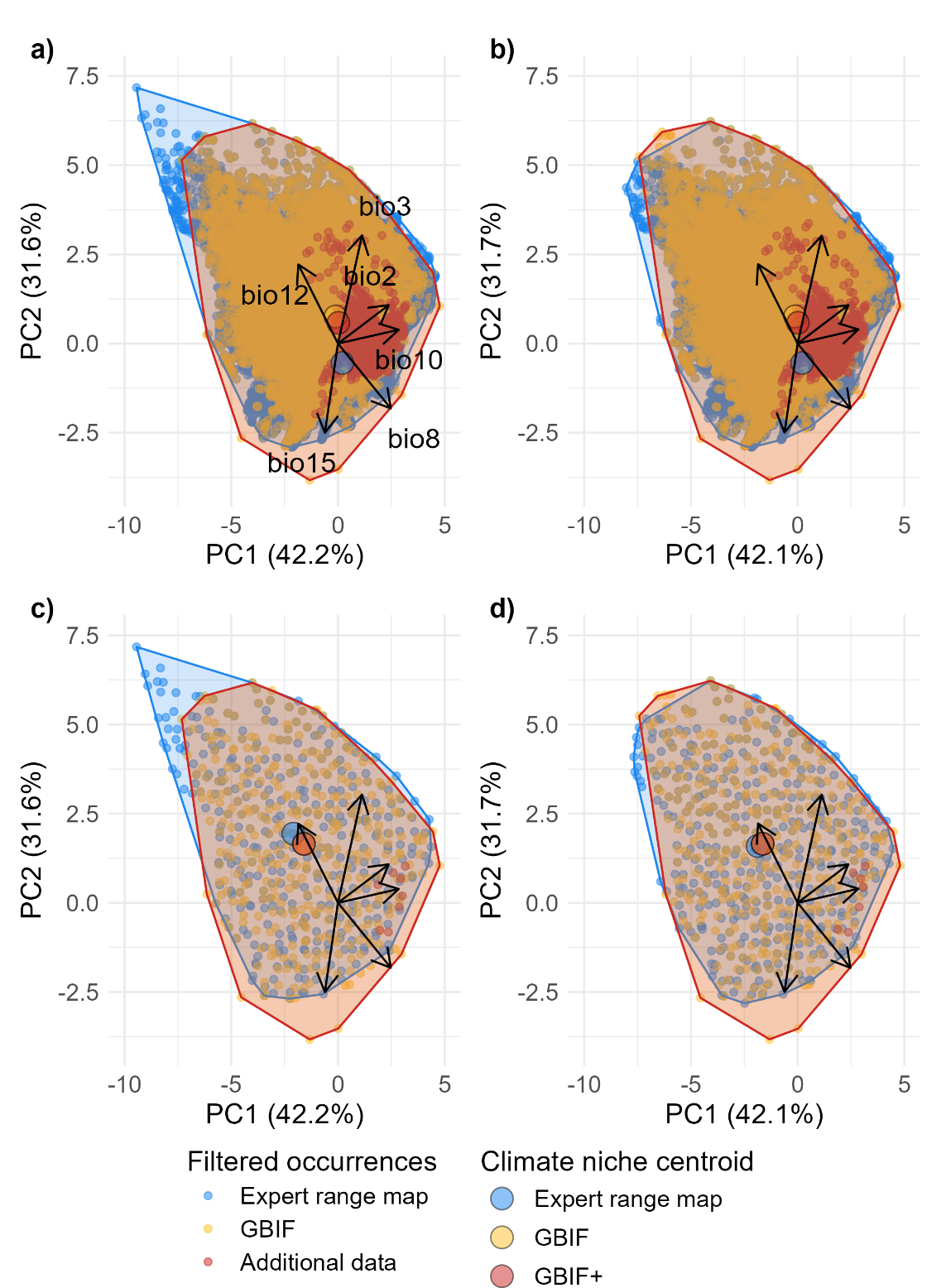

**Figure S2. Comparison of climate niches modeled with different data preparation, with an example of *Campanula rotundifolia***. a) climate niche modeling without particular data preparation; b) outlier removal of expert range map dataset (*convex hull* *peeling process*) but no environmental subsampling; c) no peeling process but environmental subsampling; and d) both peeling process and environmental subsampling. The latter option was used for the analysis.
